# Environmental carcinogens disproportionally mutate genes implicated in neurodevelopmental disorders

**DOI:** 10.1101/2021.12.17.473207

**Authors:** Brennan H. Baker, Shaoyi Zhang, Jeremy M. Simon, Sarah M. McLarnan, Wendy K. Chung, Brandon L. Pearson

**Author notes:** Author for correspondence: 722 W. 168^th^ St., New York, NY 10032.

## Abstract

*De novo* mutations contribute to a large proportion of sporadic psychiatric and developmental disorders, yet the potential role of environmental carcinogens as drivers of causal *de novo* mutations in neurodevelopmental disorders is poorly studied. We demonstrate that several mutagens, including polycyclic aromatic hydrocarbons (PAHs), disproportionately mutate genes related to neurodevelopmental disorders including autism spectrum disorders (ASD), schizophrenia, and attention deficit hyperactivity disorder (ADHD). Other disease genes including amyotrophic lateral sclerosis (ALS), Alzheimer’s disease, congenital heart disease, orofacial clefts, and coronary artery disease were generally not mutated more than expected. Our findings support a new paradigm of neurodevelopmental disease etiology driven by a contribution of environmentally induced rather than random mutations.

## Background

While cancer epidemiologic studies have a long history of integrating genetic and environmental factors into disease causation,^1^ researchers have not readily implicated environmentally-induced mutations as etiological drivers of neurodevelopmental disorders (NDD) and other diseases. *De novo* mutations contribute to a large proportion of sporadic cases of ASD, schizophrenia, and intellectual disability,^2-5^ yet the underlying mutational processes have not been interrogated, or have been attributed to intrinsic mutational processes rather than environmental carcinogens. Similarly, environmental exposures may be responsible for a large proportion of NDD.^6-8^ Genetic mutation remains a strong yet generally untested candidate mechanism that may link environmental exposures to neurodevelopment. For instance, PAHs – a class of chemicals found in tobacco smoke and air pollution – form metabolites in the body that bind with DNA and promote mutation.^9^ Consequently, PAHs are well known causes of cancer.^10-12^ Epidemiologic studies have linked prenatal PAH exposure to cognitive developmental delays, reduced intelligence, and ASD.^13-16^ However, no studies have examined whether mutations in NDD genes induced by PAHs and other environmental exposures contribute to these epidemiologic associations despite evidence that NDD genes are generally longer^17-19^ and show considerable overlap with cancer driver genes.^20,21^ To test the hypothesis that NDD genes are more susceptible to mutagens, we analyzed a whole genome sequencing (WGS) dataset containing nearly 200,000 single nucleotide substitution mutations in human induced pluripotent stem cell (hiPSC) clonal lines exposed to 12 classes of environmental carcinogens.^22^ Here, we assessed the genes susceptible to environmental mutation and the associated diseases by 1) evaluating gene ontology for top mutated genes; 2) we developed an online tool for assessing the propensity of the 12 mutagen classes to cause mutations in gene sets associated with specific human diseases (www.environmentalmutation.com). After identifying genes and diseases that are more susceptible to environmental mutation, we 3) investigated gene length, expression, GC content, and bulky adduct repair as potential drivers of elevated mutability using Quasi-Poisson models.

## Results and Discussion

We compared the observed rates of exposure-induced mutations in disease-related gene sets with the expected rates of mutations based on control genes randomly sampled from the genome. Disease gene sets contained 91 ASD,^23^ 104 schizophrenia,^24^ 25 ADHD,^25^ 33 Alzheimer’s,^26^ 18 ALS,^27^ 81 type 2 diabetes, 80 coronary artery disease, 96 obesity, 253 congenital heart disease, and 31 orofacial cleft genes (Table S1). Gene sets were either curated (i.e., in review articles or by scientific organizations) or based on genes with significant disease-associated loci from genome wide association studies (GWAS). We included adult onset, congenital, heritable, and life-style-associated diseases to determine if NDD enrichment was specific. Since our analyses were restricted to just a handful of disease gene sets and results could depend on the methods of gene set curation, we created an online tool where custom gene lists can be queried using the algorithm we generated (www.environmentalmutation.com). Gene overlap between sets was minimal (Figure S1).

To determine expected mutation rates, we randomly sampled 1,000 sets of 300 genes from the human genome and used the hiPSC mutation dataset to calculate average rates of mutation per-gene-per-treated hiPSC subclone within each exposure class. To characterize the degree to which certain disease gene sets were mutated more than expected, we compared these hypothesized expected mutation rates to the mutation rates for each disease gene set within each environmental exposure using exact binomial tests. This analysis was repeated for mutations in entire genes (Figure 1A, Table S2), as well as coding sequence (CDS) mutations (Figure 1B, Table S2). Although entire genes contain introns and other non-coding sequence, a large proportion of GWAS signals map to non-coding regions, ^28^ so variants in these loci may still contribute to disease.

**Figure 1.**
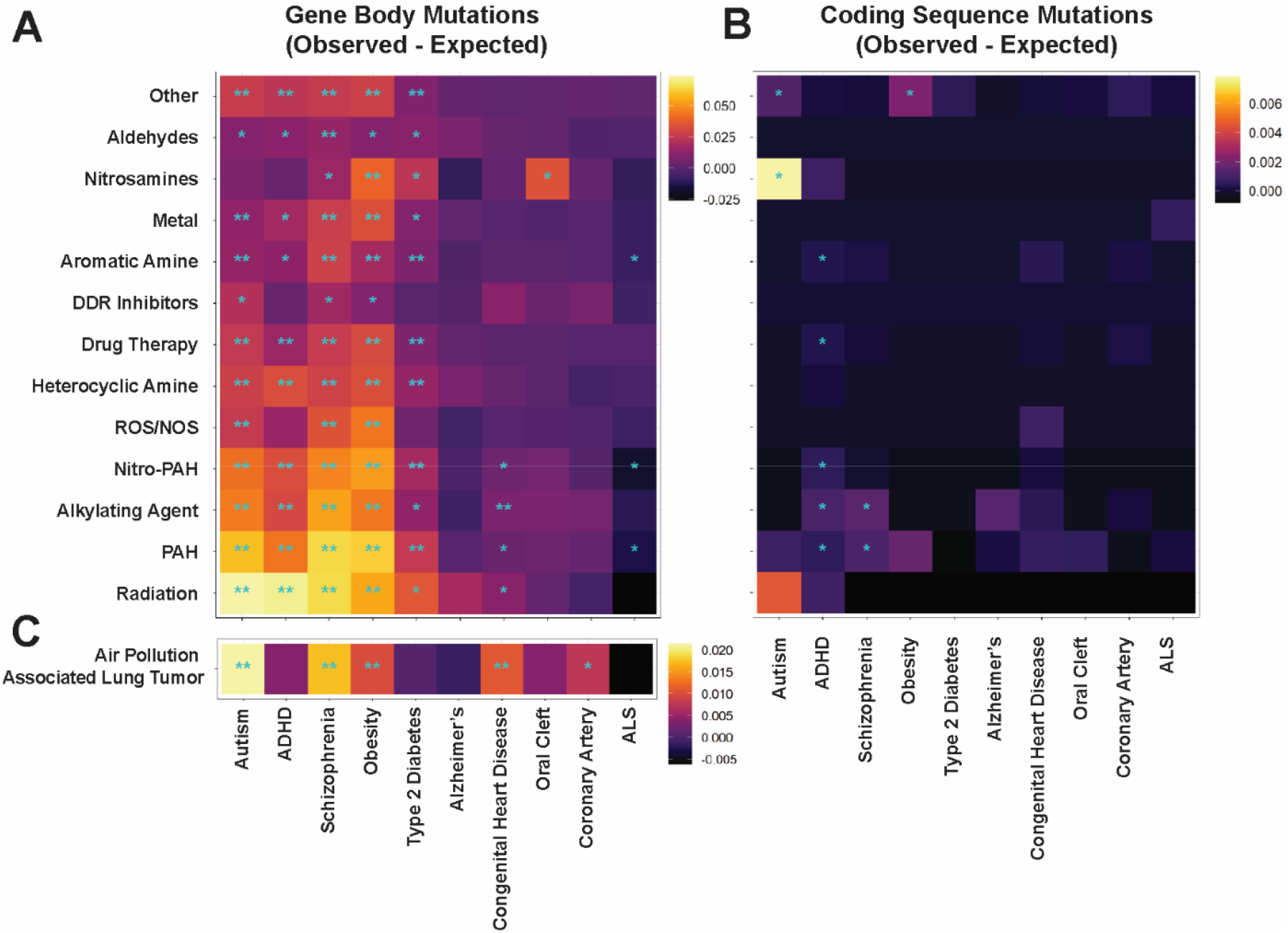
Mutation vulnerability of disease associated genes to various carcinogens. Heatmaps of observed minus expected mutations in disease gene sets for (A) mutations in hiPSC gene bodies following chemical treatment, (B) mutations in hiPSC coding sequences following chemical treatment, and (C) mutations in 14 human lung cancers from individuals living in highly polluted regions. Significant levels from exact binomial tests: * p<.05; ** Bonferroni-adjusted p<.05.

Among the nearly 200,000 substitutions, 92,204 occurred in known genes. The most mutagenic exposures were radiation and PAHs, which induced an average of 0.066 and 0.058 substitution mutations per-gene-per-treated hiPSC subclone, respectively across our 10 disease related gene sets (Figure 1A, Table S3). ASD, ADHD, schizophrenia, obesity, and type-2 diabetes genes were mutated significantly more than expected by almost every chemical class. Congenital heart disease, oral cleft, and ALS were rarely mutated more than expected; while Alzheimer’s and coronary artery disease were never mutated more than expected (Figure 1A, Table S2). (Figure 1). Among the 2,061 identified coding sequence variants, these overarching patterns were not observed. While there was evidence for exposure causing more coding sequence mutations than expected for 10 specific exposure/disease combinations, none of them remained statistically significant following Bonferroni correction (Figure 1B).

To cross-validate our approach, we repeated the gene mutation analysis in an independent dataset of human WGS data from 14 lung cancers from individuals living in highly polluted regions.^29^ Because PAHs are a major component of pollution, we hypothesized that mutational patterns would be similar between these samples and the PAH-treated hiPSCs. The greatest increases in observed over expected gene mutations were in genes related to ASD, schizophrenia, and obesity (Figure 1c). However, contrasting with the mutational patterns in PAH-exposed hiPSCs, ADHD genes were not mutated more than expected, and genes associated with congenital heart defects were mutated more than expected even after Bonferroni correction (Figure 1c).

We performed gene ontology analysis on the 692 coding sequence (CDS) variants in PAH-treated hiPSCs. Enriched gene ontologies were closely related to neurodevelopment: neuron projection, neuron part, calcium ion binding, and cell projection part (Table S4). Furthermore, the enriched plasma membrane term may be related to metabolic diseases including obesity and type 2 diabetes.^30^ Similar results were obtained when the analysis included all 2,061 CDS mutations across all hiPSCs exposed to environmental mutagens. For instance, the top three gene ontology terms were neuron projection guidance, sensory organ morphogenesis, and cell morphogenesis involved in neuron differentiation (Table S5), supporting our hypothesis that NDD genes are particularly vulnerable to environmental mutagens.

ASD-implicated NDD genes tend to be longer than other genes.^17,31^ To examine if vulnerability of neurodevelopmental genes or CDS to mutagens is attributable to sequence length, we plotted the distributions of entire gene and CDS lengths for our disease gene sets, along with the distributions of lengths for entire genes and CDS mutated entirely at random (Figure 2A, E). Random mutations were modeled by randomly sampling (i.e., mutating) 100,000 nucleotides from all genes or all CDS in the human genome, so the probability of a sequence being mutated was entirely governed by its length. We additionally modeled the effects of sequence length, expression, and percent GC content on mutation rate using multivariate Quasi-Poisson regressions, with separate models for mutations in CDS and entire genes (Figure 2).

**Figure 2.**
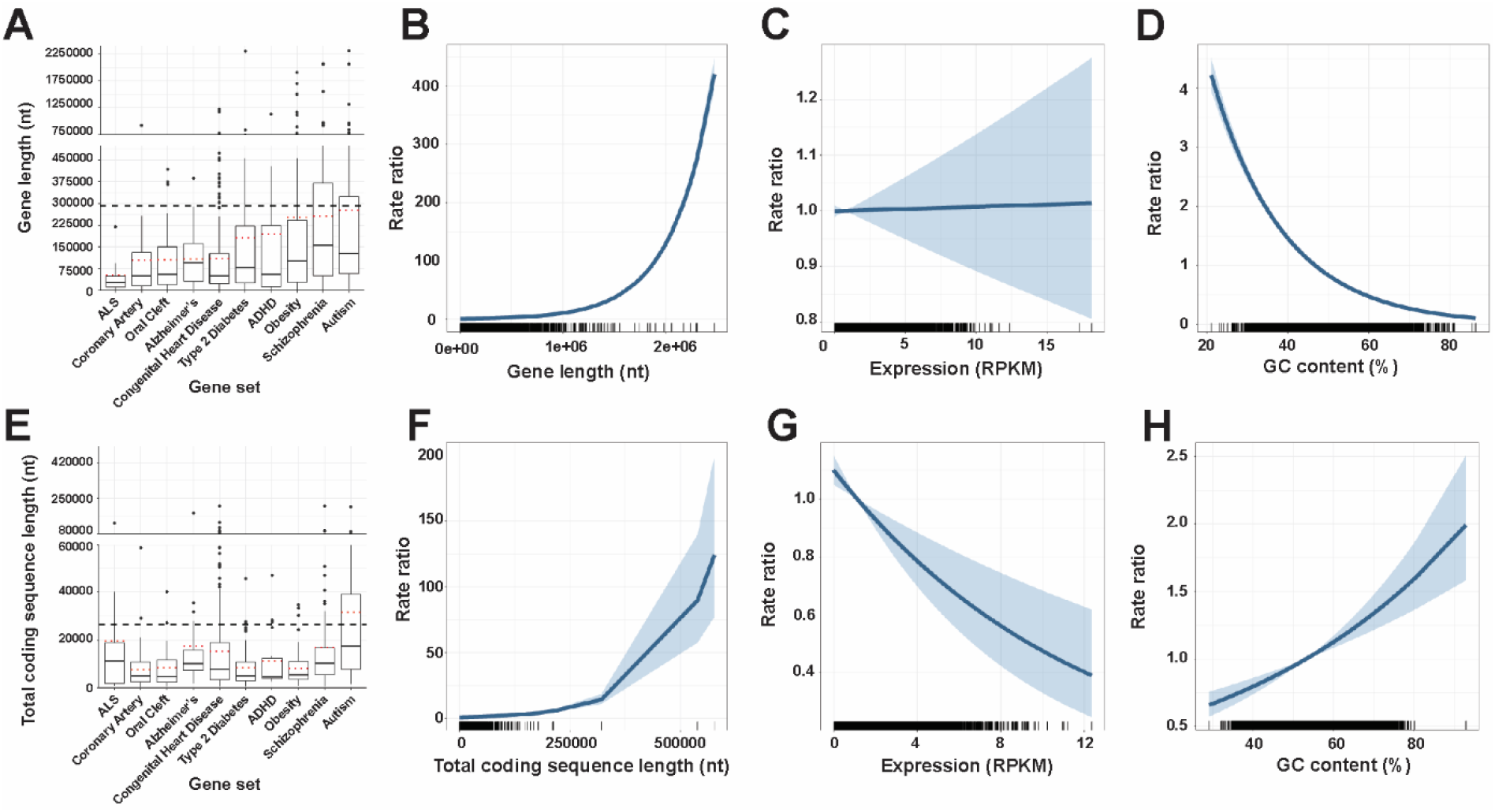
Gene length and other determinants of mutation vulnerability across gene bodies (A-D) and coding sequences (E-H). Gene length boxplots with 1.5 interquartile range (IQR) whiskers shown for each disease implicated gene set (A). Red dotted lines indicate mean gene length, and dashed black line indicates mean gene length of randomly mutated genes. Longer gene length is associated with higher mutation frequencies across carcinogen treated cells (B). Expression is not associated with altered mutation risk across gene bodies (C). Higher GC content is associated with decreased mutation risk across gene bodies (D). Coding sequence (CDS) length boxplots with 1.5 IQR whiskers shown for each disease implicated gene set (E). Longer coding sequence is associated with a modestly increased risk of mutation (F). Higher gene expression is associated with reduced CDS mutations (G). In contrast to the gene body, higher GC content is associated with increased risk of CDS mutation (H).

Average gene lengths of disease gene sets were consistent with patterns of mutability (Figure 2A). For instance, ASD, schizophrenia, obesity, and ADHD genes, which were on average longer than other disease-related genes, had the greatest increases in observed versus expected chemical-induced mutations, while ALS, coronary artery disease, oral cleft, and Alzheimer’s disease genes, which are much shorter in length, were not mutated more than expected (cf. Figure 1A, Figure 2A). Genes mutated at random were on average longer than all disease gene set genes, further indicating that gene length was a strong driver of mutability. On the other hand, average CDS lengths for most disease gene sets were similar to each other (Figure 2E).

Quasi-Poisson regressions further supported a stronger role of sequence length in gene but not CDS mutation number. Controlling for expression and GC content, each interquartile range increase (IQR) in gene length was associated with a 1.104-fold increase in gene mutation rate (rate ratio (RR) = 1.104, 95% CI [1.102, 1.105]), while an IQR-increase in CDS length was associated with a 1.050-fold increased CDS mutation rate (RR = 1.050, 95% CI [1.045, 1.055]) (Figure 2B,F).

Quasi-Poisson regressions also showed significant effects of GC content and expression on gene and CDS mutability. In this analysis, expression was not associated with mutability for genes (RR = 1.001, 95% CI [0.988, 1.014]), while each IQR increase in expression was associated with a 13% decreased mutation rate for CDS (RR = 0.870, 95% CI [0.812, 0.931]; Figure 2C,G). This corroborates prior work showing that lowly expressed genes harbor more mutations,^32^ which may be driven by transcription-coupled DNA repair.^33,34^ Similarly, the effect of GC content was different for genes and CDS. Each IQR-increase in gene GC content was associated with a 0.524-fold decreased mutation rate (RR = 0.524, 95% CI [0.508, 0.539]), while each IQR-increase in CDS GC content was associated with a 1.274-fold increased mutation rate (RR = 1.274, 95% CI [1.175, 1.381]) (Figure 2D,H). These results are consistent with prior studies indicating that the effect of GC content on mutation rate varies over different genomic scales. GC content across entire genes may reflect higher order DNA structure, and increased GC content has been shown to correlate with decreased mutation rate at higher genomic scales.^35,36^ CDS GC content, however, may more accurately reflect the effect of local GC content on mutability. Cytosines may experience higher mutation rates than other bases because methylated cytosines in CpG dinucleotides are vulnerable to deamination into thymidine. Furthermore, these mutations occur at higher rates in regions with higher local GC content.^37^

To further explore the role of local sequence context on mutability, we aligned 7-mers centered on each gene or CDS mutation, along with 50,000 7-mers randomly sampled from the human genome (Figure S2). Mutated regions were GC enriched, while randomly sampled 7-mers contained equal proportions of each nucleotide. G and C were more highly enriched in 7-mers centered on CDS mutations compared to 7-mers centered on gene mutations. Thus, CDS mutations may be governed more strongly by local GC content. When aligning 7-mers on each gene or CDS mutation stratified by chemical exposure, this pattern held for some but not all chemical classes (Figure S3).

We also examined the role of local sequence context by generating COSMIC signatures^38^ for *de novo* mutations in individuals with neuropsychiatric diseases, including approximately 50,000 ASD cases,^39^ 617 schizophrenia cases,^3^ and 145 individuals with severe intellectual disability.^2,40^ Single base substitution enrichments for all neuropsychiatric cases and controls were clock-like/aging associated signatures (i.e. SBS1, Figure S4), which are enriched for NpCpG to NpTpG substitutions. The SBS1 signature does not resemble any of the chemical mutation signatures identified by Kucab and colleagues.^22^ This preliminary analysis suggests that *de novo* mutations in ASD, schizophrenia, and intellectual disability may reflect sporadic mutational processes rather than chemical-induced mutation. However, it is also possible that mutational signatures generated from hiPSC cultures are not readily comparable to *in vivo* human mutational signatures. For instance, methylated CpG sequences are disproportionately targeted by environmental carcinogens such as PAHs, which form guanine adducts that induce G to T transversions at methylated CpGs.^41^

Finally, to determine if NDD genes are linked to PAH adduct repair, we analyzed an existing genome-wide PAH adduct repair assay dataset^42^ to see if adducts are preferentially located in specific disease-related gene sets. We computed the average DNA damage enrichment across each gene or CDS in all 10 disease-related gene sets using the deepTools2 ‘computeMatrix’ function.^43^ By default, the ‘computeMatrix’ function scales input sequences to the same length. We used one-way ANOVA to compare levels of DNA damage in genes, which were normally distributed, between disease gene sets, and performed pairwise contrasts with a false discovery rate correction. Kruskal-Wallis and Dunn’s Test were employed for CDS DNA damage data, which were not normally distributed.

Pairwise contrasts revealed that DNA damage following PAH treatment was significantly enriched in genes of ASD-related genes compared to genes associated with coronary artery disease, congenital heart defects, and orofacial cleft (Figure S5A, Table S6). Schizophrenia genes similarly demonstrate more DNA damage compared to coronary artery disease and congenital heart defect genes. However, this pattern of increased DNA damage in neurodevelopmental diseases was not observed for CDS. In fact, ASD and schizophrenia CDS were among the diseases with the lowest levels of DNA damage (Figure S5B). Although over half of the Dunn’s Tests contrasts were significant (Table S6), differences in the mean and median CDS DNA damage enrichments between diseases were minimal (Figure S5B).

## Conclusions

We have shown that environmental carcinogens may disproportionately mutate neurodevelopmental genes. Additionally, metabolic genes such as those associated with obesity may also have increased susceptibility to environmental carcinogens. Genes mutated by PAHs were overwhelmingly enriched for neurodevelopmental processes. Neurodevelopmental genes may be particularly sensitive to mutation because the transcriptome of neural tissues, especially neurons, is biased toward longer genes.^17,31^ While gene length and gene mutation number were highly correlated, other factors like local nucleotide composition may govern the mutability of protein coding sequences. The mutational signatures of these neuropsychiatric disorders closely mirrored the most common mutational signatures in cancer, complementing known overlaps between developmental disease and cancer genes. Environmentally induced mutations may play a greater role in neurodevelopmental disease than previously assumed. Rather than attributing sporadic neurodevelopmental diseases to intrinsic mutational processes, this work suggests that a proportion of genetic neurodevelopmental disease risk may be explained by environmental mutagenesis.

## Methods

We analyzed the substitution mutations from 324 hiPSC subclones dosed with 79 environmental carcinogens.^22^ From whole-genome-sequencing data at ∼30-fold depth, Kucab and colleagues (2019) called mutations in subclones subtracting on the primary hiPSC parental clone. The ensembl variant effect predictor^44^ was used to predict the consequence of each mutation from its genomic location and type. Gene ontology analysis was performed in FUMA with ensembl version 92 and all protein coding genes set as the background.^45^

Gene and CDS start and stop positions were obtained from GENCODE Release 38 (https://www.gencodegenes.org/human/) and used to calculate genomic sequence lengths (end minus start position) and GC content (proportion of sequence positions with either a G or C nucleotide). When modeling associations of sequence properties with CDS mutations, CDS lengths and GC content were computed per gene: all CDS segments within each gene were summed as the total coding sequence length, and CDS GC content was calculated per gene rather than per CDS segment. Gene expression data were reads per kilobase of transcript per million mapped reads (RPKM) obtained from RNA-seq of hiPSCs generated using the Sendai virus method,^46^ the same method used to create the hiPSCs used by Kucab and colleagues (2019). After excluding genes with missing length, expression, or GC content data, Quasi-Poisson models included 75,756 gene and 1,852 CDS mutations. Analyses were performed with the statistical package R.^47^

Genome-wide PAH adduct repair data come from translesion excision repair-sequencing (tXR-seq) of GM12878 cells, which were grown to ∼80% confluence before treatment with 2 μM benzo[a]pyrene diol epoxide-deoxyguanosine for 1 h at 37 °C in a 5% CO_2_ humidified chamber. tXR-seq captures all DNA damage, regardless of whether or not it is repaired.^42^

## Declarations

Data reflect secondary data analysis of published data; it was therefore exempt of IRB approval. Data are available from the primary sources cited. The SPARK gene variants are available to approved researchers through SFARI upon review. Authors declare no competing interests. Support for this work was provided by the following NIH grants: P30ES009089, R21ES032913, and R24ES029489.

## Acknowledgements

Dr. Barbara Corneo of the Columbia Stem Cell Initiative provided valuable intellectual contribution to the conception of our study.

