## Supplementary figures and images for "Environmental carcinogens disproportionally mutate genes implicated in neurodevelopmental disorders"

### Figure S

#### Slide 1
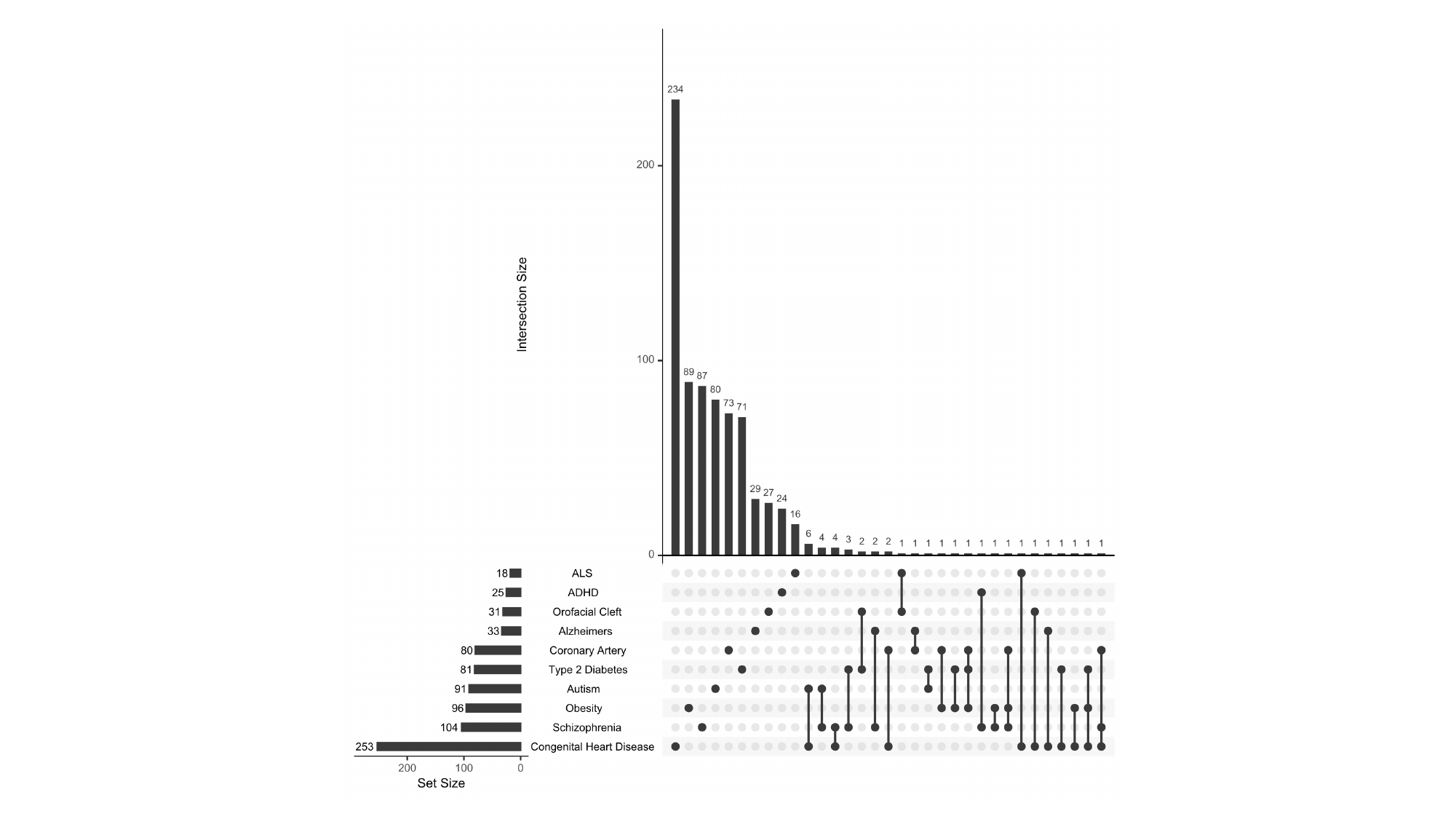

#### Slide 2
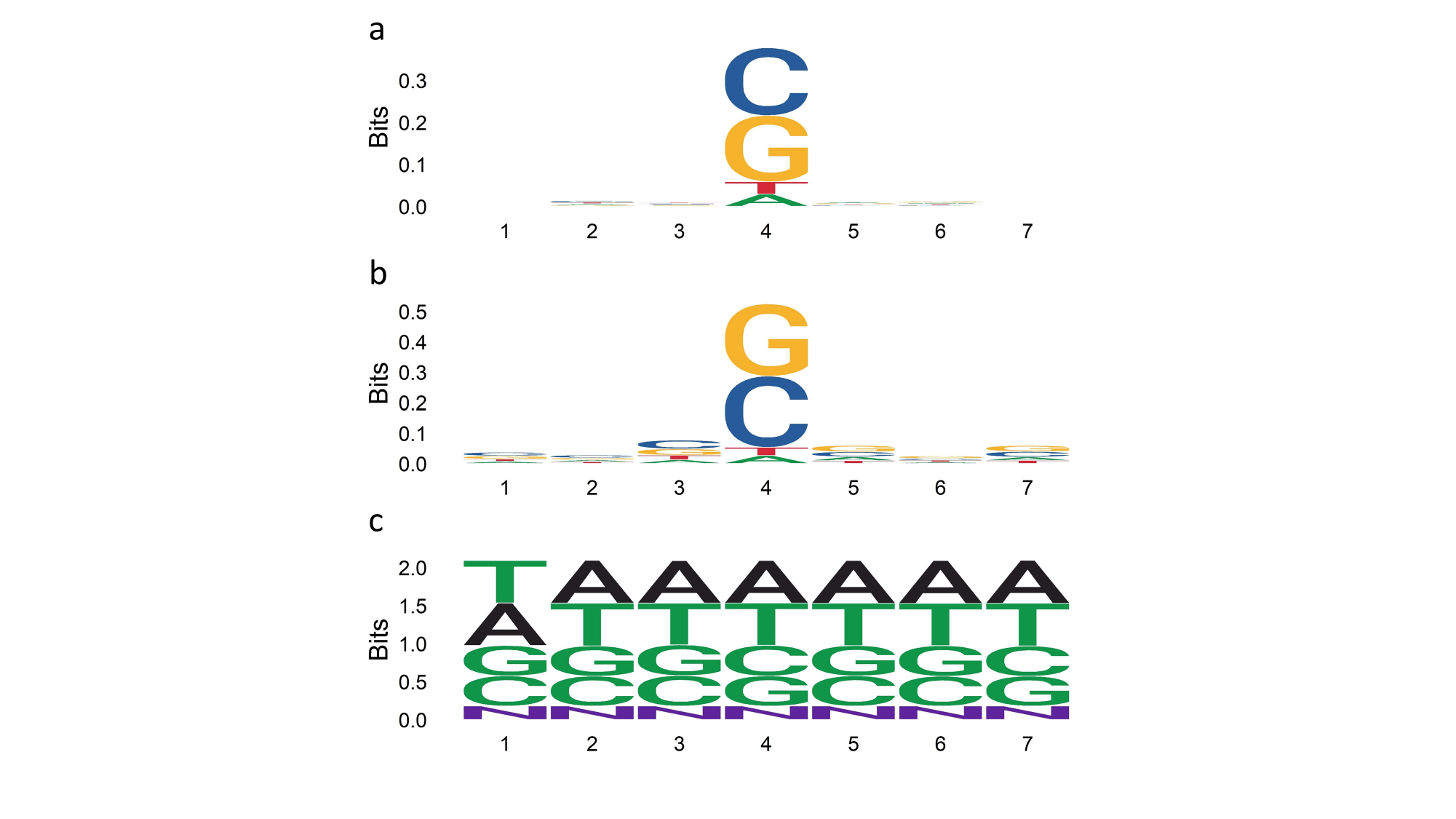

#### Slide 3
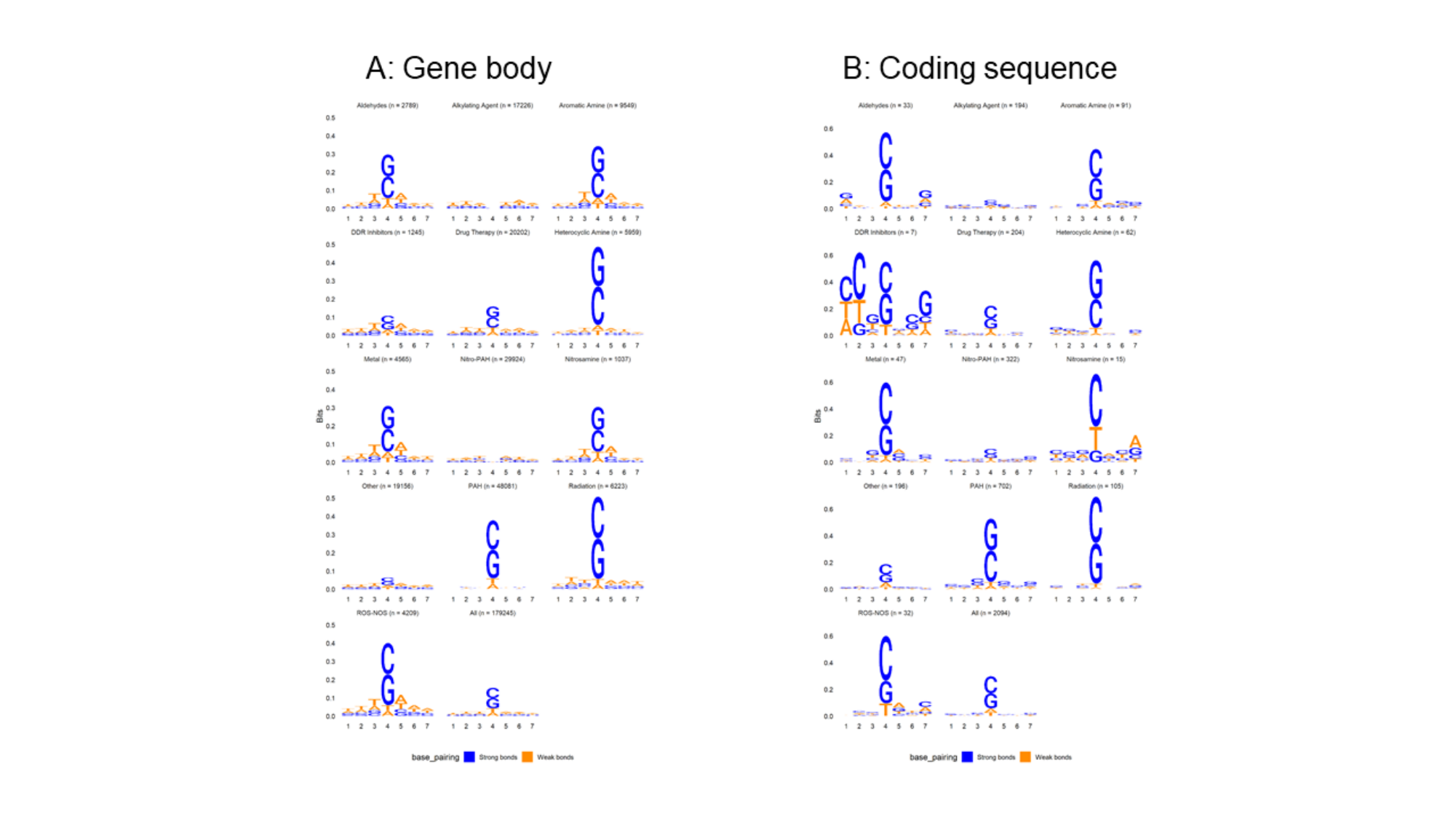

#### Slide 4
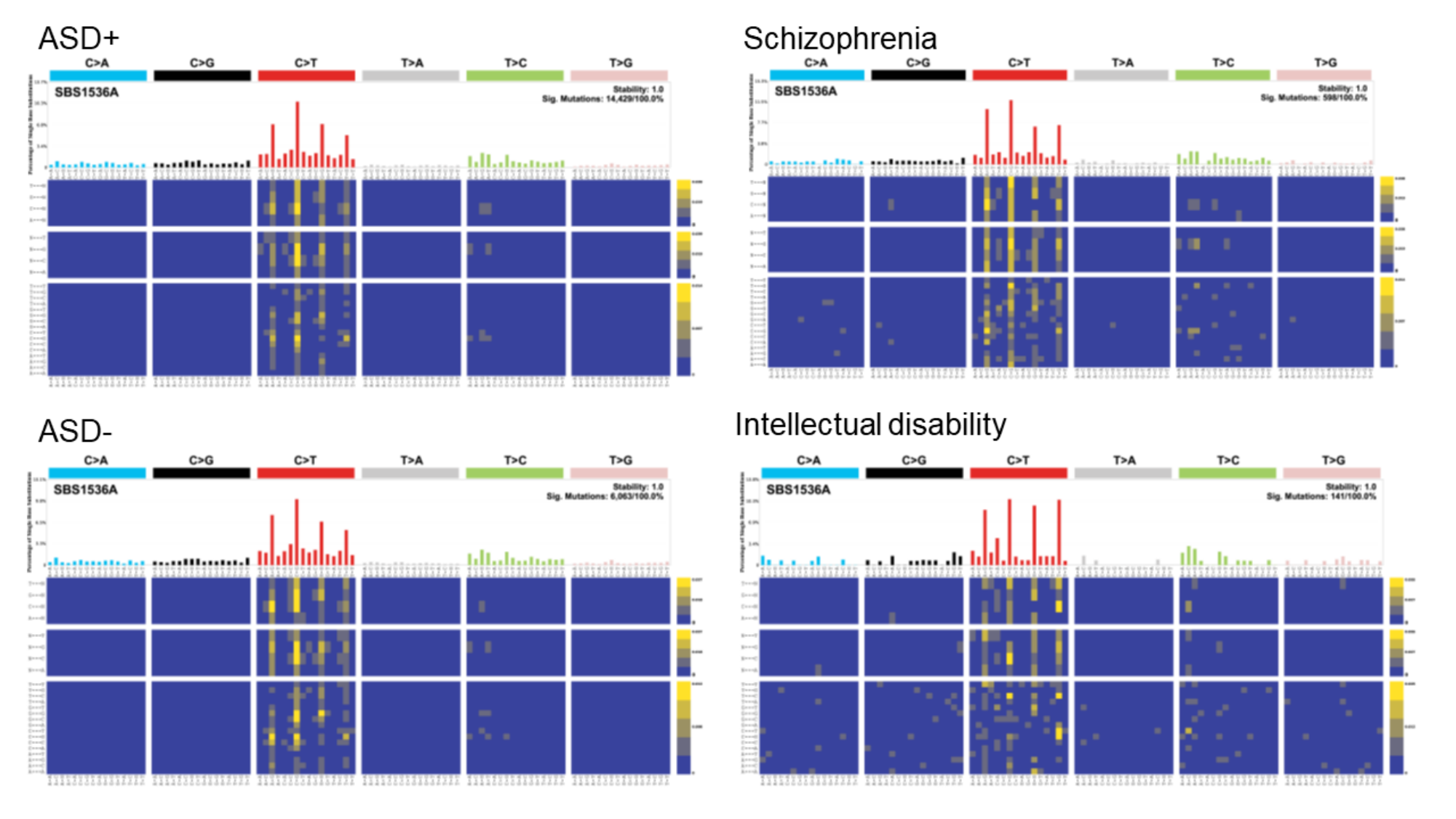

#### Slide 5
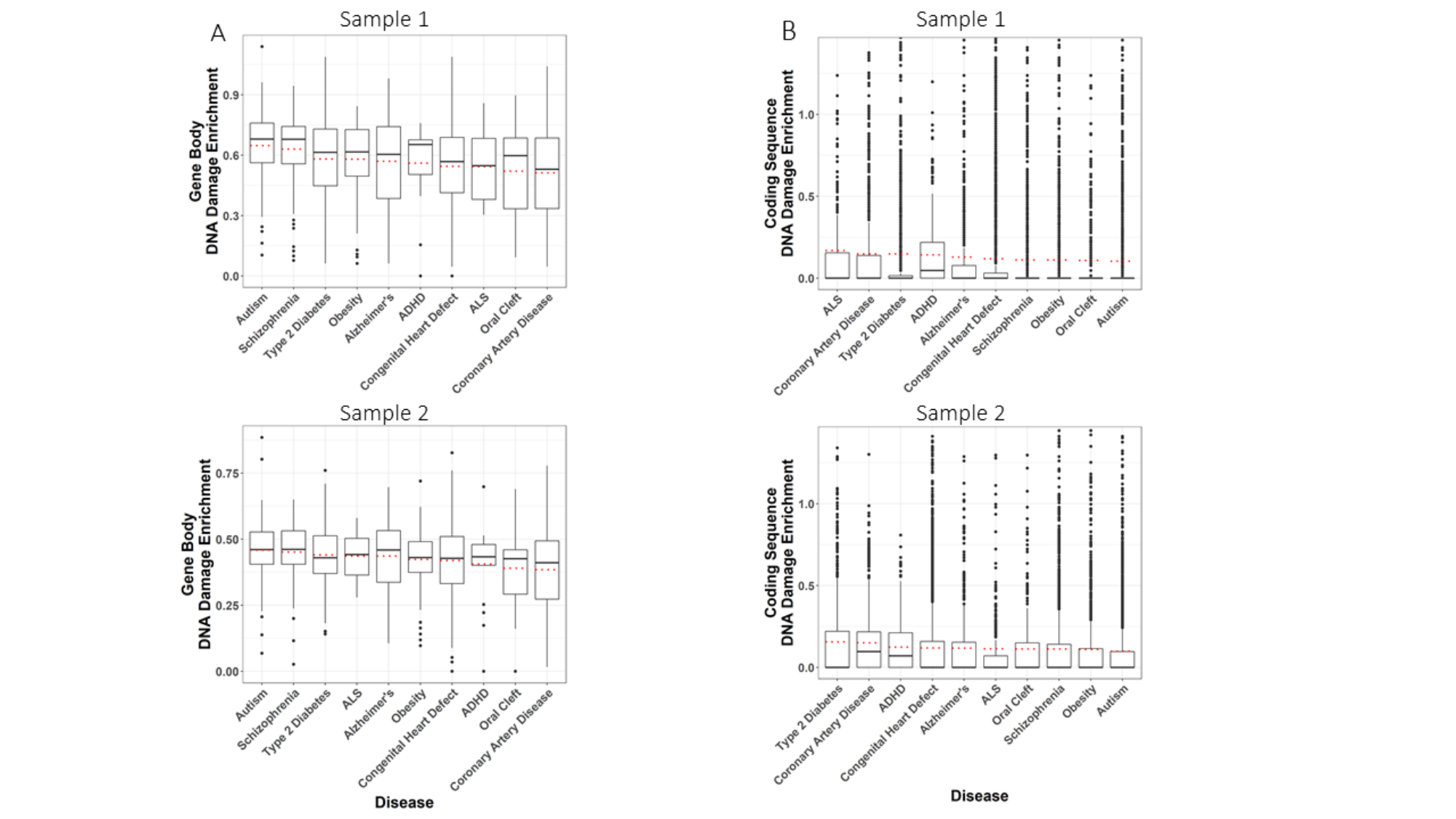
